# Continuous force measurements reveal no inhibitory control deficits in Parkinson’s disease

**DOI:** 10.1101/809780

**Authors:** Jade S. Pickering, Jennifer McBride, Iracema Leroi, Ellen Poliakoff

## Abstract

Suppression of unwanted motor responses can be disrupted by Parkinson’s disease. People with Parkinson’s (PwP) can show maladaptive reward-driven behaviours in the form of impulse control behaviours, which are associated with use of the dopaminergic treatments used to alleviate the motor symptoms of the disease. However, the effects of Parkinson’s itself on impulsive behaviour and control are unclear – empirical studies have yielded mixed findings, and some imaging studies have shown a functional deficit in the absence of a measurable change in behaviour. Here, we investigated the effects of Parkinson’s on response activation and control by studying the dynamics of response in standard inhibitory control tasks – the Stop Signal and Simon tasks – using a continuous measure of response force. Our results are largely in favour of the conclusion that response inhibition appears to be intact in PwP, even when using a more sensitive measure of behavioural control relative to traditional button-press measures. Our findings provide some clarity as to the effects of Parkinson’s on response inhibition and show continuous response force measurement can provide a sensitive means of detecting erroneous response activity in PwP, which could also be generalised to studying related processes in other populations.

**Open data:** Data and analysis can be found on the Open Science Framework (osf.io/kx6h3/)

**Pre-print template:** Wiernik, B.M. (2019, October 11). Preprint templates. osf.io/hsv6a/

---

Parkinson’s disease is a neurodegenerative disorder affecting around 1% of all adults over the age of Jade S. Pickering, D Jennifer McBride, D Iracema Leroi, Ellen Poliakoff, 60 (Tysnes & Storstein, 2017). Parkinson’s is associated with significant loss of dopaminergic cells in the substantia nigra pars compacta, which in turn supplies dopamine to the dorsal striatum of the basal ganglia (Dauer & Przedborski, 2003) and frontal regions (Jahanshahi et al., 2015). This neural loss in Parkinson’s has a profound effect on the motor system: people with Parkinson’s (PwP) can experience muscle rigidity, tremor, freezing of gait, and slowness of movement (bradykinesia; Jankovic, 2008). In addition to PwP being slow to initiate and execute movements, they can also have difficulty with the *inhibition* of prepotent responses (e.g., Gauggel, Rieger, & Feghoff, 2004; Nombela, Rittman, Robbins, & Rowe, 2014). Sometimes, deficits in inhibition and control can manifest as impulse control behaviours (ICBs), including pathological gambling, hypersexuality, binge eating, and compulsive shopping (Voon, 2015). Recent estimates suggest that up to 50% of PwP develop an ICB (Corvol et al., 2018), which can negatively impact on quality of life (Leroi et al., 2011; Phu et al., 2014).

However, “impulsivity” is a complex and multifaceted construct; Antonelli et al. (2011) distinguished between cognitive impulsivity – which is characterized by altered decision-making (e.g. risk-taking, altered time-perception, and avoidance of waiting), and motor impulsivity – which is associated with a relative inability to inhibit prepotent responses. Response conflict and inhibition have been widely studied experimentally using a variety of tasks, including the Go/No-Go (e.g. Gomez et al., 2007), Stop Signal (Verbruggen & Logan, 2008), and Simon tasks (Simon, 1967, 1990). In the Go/No-Go task participants must respond to the presence of a Go signal on most trials (“Go” trials) but withhold their response when presented with the No-Go signal on a small number of trials. Commission errors are the primary measure of interest; instances where participants fail to withhold their response on No-Go trials. In the related Stop Signal task, participants must respond as quickly as possible to a Go stimulus on each trial but withhold that response when this Go signal is followed by a Stop signal (presented on a minority of trials). Researchers typically calculate the stop signal reaction time (SSRT) –an estimate of the time needed to successfully inhibit a response which has already been initiated. Thus, the Stop Signal task requires *cancellation* of an in-progress response, whereas the Go/No-go task requires participants to *withhold* a prepotent response.

In contrast, the Simon task (Simon, 1967, 1990) measures inhibitory control over competing motor responses. For example, a typical set-up might include instructions to the participant to respond with the left button when they see a yellow stimulus, and the right button when they see a blue stimulus. Crucially, the stimulus may appear on the left or the right of the screen, but the location of the stimulus is not relevant to the participant’s task (which is to respond according to stimulus colour). Therefore, the stimulus’s location might prime a response that is congruent (same side) or incongruent (opposite side) with the response required by the task instructions On incongruent trials, the automatically activated response elicited by the location of the stimulus must be inhibited in favour of the goal-directed response according to stimulus colour (or another visual feature), which results in longer response times (RTs) and reduced accuracy for incongruent compared to congruent trials. Therefore, the Simon task measures resolution of conflict between competing motor responses which have been simultaneously activated by different aspects of the stimulus.

Although Parkinson’s has been associated with disrupted inhibitory control and a high incidence of ICBs, empirical studies investigating the effects of Parkinson’s on response conflict and inhibition have produced mixed findings. For example, some studies using the Simon task have found that PwP show greater interference between competing responses (the difference in RTs for incongruent versus congruent trials e.g., Houvenaghel et al., 2016; van Wouwe et al., 2016) compared to healthy controls (HCs), whereas others have found no significant group differences (Wylie, Ridderinkhof, Bashore et al., 2010; Wylie, Ridderinkhof, Elias et al., 2010). Moreover, whilst some studies have shown that PwP produce more commission errors on the Go/No-Go task compared to HCs (Geffe et al., 2016; Nombela et al., 2014), others have reported no group differences (de Rezende Costa et al., 2016; Georgiev, Dirnberger, Wilkinson, Limousin, & Jahanshahi, 2016). A hybrid Go/No-Go task that incorporated congruent and incongruent conditions (as in a Simon task) showed a larger interference effect for PwP relative to healthy controls in some conditions (Beste, Dziobek, Hielscher, Willemssen, & Falkenstein, 2009). Similarly, there is some evidence to suggest that PwP have longer SSRTs compared to HCs (and therefore reduced inhibitory control e.g., Gauggel et al., 2004; Nombela et al., 2014), whereas others have found no difference (Bissett et al., 2015). Still further studies have shown a functional deficit in PwP (e.g. differences in the blood oxygenation level-dependent (BOLD) signal in the fronto-striatal-thalamic loop during the Go/No-Go task, and the inferior frontal gyrus in the Stop Signal task) relative to HCs, even in the absence of an observable behavioural deficit (e.g. Baglio et al., 2011; Vriend et al., 2015).

Thus, it remains unclear whether or how Parkinson’s may affect control over actions. However, there are substantial differences between studies – in terms of task, methods, analysis, and participants – which make it difficult to draw clear conclusions. For example, most studies investigating motor activation and/or control compare the time taken to respond in different conditions and report an overall central tendency for each condition. However, such a measure of central tendency does not elucidate differences in higher-order characteristics of the RT distribution and can be skewed by variability between participants (Ratcliff, 1993). More recently, some researchers have been comparing performance on tasks or conditions across the whole RT distribution. When applied to tasks measuring inhibition or conflict, these distributional analyses aim to temporally dissociate impulsive errors at the fast end of the RT distribution from failed inhibition at the slow end. According to the activation suppression model (Ridderinkhof, 2002a, 2002b; van den Wildenberg et al., 2010) slower RTs allow more time for selective suppression of the automatic response to build up, whereas faster RTs do not allow sufficient time for inhibition and can result in fast, impulsive errors. This is visible by plotting accuracy (in conditional accuracy functions) or the RT interference effect (in delta plots) as a function of RT (see van den Wildenberg et al., 2010 for a review). Using these methods, studies have consistently revealed that PwP show deficits in successful inhibition of responses at the slow end of the RT distribution (van Wouwe et al., 2016; Wylie, Ridderinkhof, Bashore et al., 2010; Wylie, Ridderinkhof, Elias et al., 2010), but are no more susceptible to fast impulsive errors than HCs on the Simon task.

Moreover, many studies infer response inhibition and conflict by comparing the time it takes participants to press a button in response to different stimuli. However, button-press measures do not capture the process of response preparation, competition, and control. The binary nature of button press measures means that either a button press is detected, or it is not, and small amounts of force which are applied to a button (and reflect ongoing cognitive control) might escape detection. The tools that have been used to measure these processes are not ideally suited to the task, and thus might contribute to the unclear nature of the effects of Parkinson’s disease on inhibitory control. The findings of Baglio et al. (2011) and Vriend et al. (2015) suggest that there is a need for a more sensitive behavioural measure to examine response inhibition in Parkinson’s. An alternative method of response measurement, therefore, is to directly measure response force. Indeed, such measures have been used successfully to measure simultaneous activation of competing motor plans, inhibition, and control in healthy adult participants (McBride, Sumner, & Husain, 2012, 2018) as well as neurological patients (McBride, Sumner, Jackson, Bajaj, & Husain, 2013), and similar measures have provided important constraints on computational models of human behaviour (Servant, White, Montagnini, & Burle, 2015).

In the present study we sought to examine the effects of Parkinson’s disease on response inhibition and control by having the *same* participants complete two different tasks measuring different kinds of inhibitory control (the SST and the Simon task), while using a sensitive measure of continuous response force. Together, this provides an opportunity to elucidate the effects of Parkinson’s disease on the dynamics of response inhibition and control.

## Materials and Methods

### Participants

25 participants (17 males, mean age 63.84 ± 5.35) with mild to moderate idiopathic Parkinson’s^1^ (Hoehn & Yahr stages 1-3) and 23 healthy control participants (12 males, mean age 68.91 ± 5.62) took part in the study (Table 1). No participants reported a history of neurological conditions (except Parkinson’s).

**Table 1.** Characteristics of the Parkinson’s and control groups^4^. Data represent ratios or means and standard deviations

|  | PwP (*n* = 25) | HCs (*n* = 25) | Statistical test |
| --- | --- | --- | --- |
| Age (years) | 63.84 (5.35) | 68.91 (5.62) | $t(46) = 3.20, p = .002^*$ |
| Education (years) | 15.72 (3.16) | 16.57 (3.34) | $t(46) = .90, p = .37$ |
| Male:Female | 17:8 | 12:11 | $\chi^2(1, N = 48) = 1.26, p = .27$ |
| Addenbrooke's Cognitive Examination | 95.04 (3.81) | 97 (2.20) | $t(38.9) = 2.20, p = .03^*$ |
| Barratt Impulsiveness Scale (total score) | 57.89 (9.54) | 51.24 (7.17) | $t(46) = 2.71, p = .009^*$ |
| BIS (attentional) | 16.42 (3.01) | 14.85 (2.90) | $t(46) = 1.83, p = .07$ |
| BIS (motor) | 21.23 (3.78) | 18.50 (2.39) | $t(46) = 2.97, p = .004^*$ |
| BIS (non-planning) | 20.52 (4.84) | 18.45 (4.20) | $t(46) = 1.57, p = .12$ |
| Handedness (L:R) | 2:23 | 2:21 | $\chi^2(1, N = 46) = .008, p = .93$ |
| Test of Premorbid Functioning | 57.56 (11.23) | 61.87 (8.23) | $t(43.88) = 1.53, p = .13$ |
| Geriatric Depression Scale | 7.37 (5.91) | 4.13 (4.09) | $t(42.83) = 2.22, p = .03^*$ |
| Disease duration (years) | 8.08 (4.53) |  |  |
| Symptom laterality (L:R) | 16:9 |  |  |
| MDS-UPDRS III | 26.64 (12.61) |  |  |
| H&Y Stage | 2 (.65) |  |  |
| Subtype (TD:PIGD)<sup>5</sup> | 8:17 |  |  |

Note: MDS-UPDRS = Movement Disorders Society Unified Parkinson's Disease Rating Scale, H&Y Stage = Hoehn & Yahr Stage, TD = Tremor dominant symptoms, PIGD = Postural instability/gait dominant symptoms. \* denotes significant differences between groups with a two-tailed alpha level of .05
<sup>4</sup> As suggested by an anonymous reviewer, we repeated the main analyses in an exploratory manner with age and GDS scores as covariates to check that the significant group differences on these measures did not affect the pattern of results. We found that our conclusions remained the same.
<sup>5</sup> The MDS-UPDRS was used to identify tremor dominant and postural instability and gait dominant patients using the same method reported by Stebbins et al. (2013).

Two patients were not receiving dopaminergic treatment during the study, 21 were taking levodopa medication, 12 were taking dopamine agonists^2^, and 18 were taking monoamine oxidase inhibitors. No patients had received deep brain stimulation. PwP were tested ON medication and had a mean score of 26.64 (± 12.61) on the Movement Disorders Society Unified Parkinson’s Disease Rating Scale Motor Section III (Goetz et al., 2008) and 2 (± .65) on the Hoehn and Yahr (1967) staging of symptom severity. All participants completed the Addenbrooke’s Cognitive Examination (Mioshi, Dawson, Mitchell, Arnold, & Hodges, 2006) to exclude significant cognitive impairment (none were excluded on this basis), the Test of Pre-morbid Functioning (Wechsler, 2011), Edinburgh Handedness Inventory (Oldfield, 1971), Geriatric Depression Scale (Yesavage et al., 1983), and the Barratt Impulsiveness Scale^3^ (Barratt, 1959). Missing data on the Geriatric Depression Scale were replaced with the total mean score for that participant.

The study was approved by an NHS Research Ethics Committee (NRES Committee North West – Liverpool Central) and was conducted in accordance with the Declaration of Helsinki.

### Tasks and Procedures

Participants performed both tasks in a darkened room and provided button press responses using a standard QWERTY keyboard that had force sensing resistors (FSRs; Interlink Electronics FSRTM 400) placed upon the A and L keys. Force data were recorded at 1000Hz and digitized using a LabJack U3 HC data acquisition device with DAQFactory Express software (version 16.2, Azeo Tech Inc.). Participants were instructed to keep the index fingers of each hand on the FSRs throughout each task so that a continuous force measurement could be recorded. Voltage change from the FSRs provided a continuous measure of response force, simultaneously and independently from the left and right hands.

#### Simon task

The Simon task was programmed in E-Prime (version 1.2, http://www.pstnet.com) and run on a computer with a flat 20inch screen (resolution of 1024×768 pixels, 75Hz refresh rate). Although actual timings were dependent on the refresh rate, the timings reported here were as programmed in E-Prime. Each trial began with a centrally presented white fixation cross (77px) on a black screen for 500-1000ms (drawn from a rectangular distribution randomly and independently on each trial). A blue or yellow circle (176px diameter) was presented at one of three locations (left, right, or centrally; that is, horizontally centred at 25%, 50%, or 75% of the screen width) (Fig. 1A). Participants were instructed to respond according to the colour of the circle as quickly and as accurately as possible, and to ignore its location on the screen.

**Fig 1.**
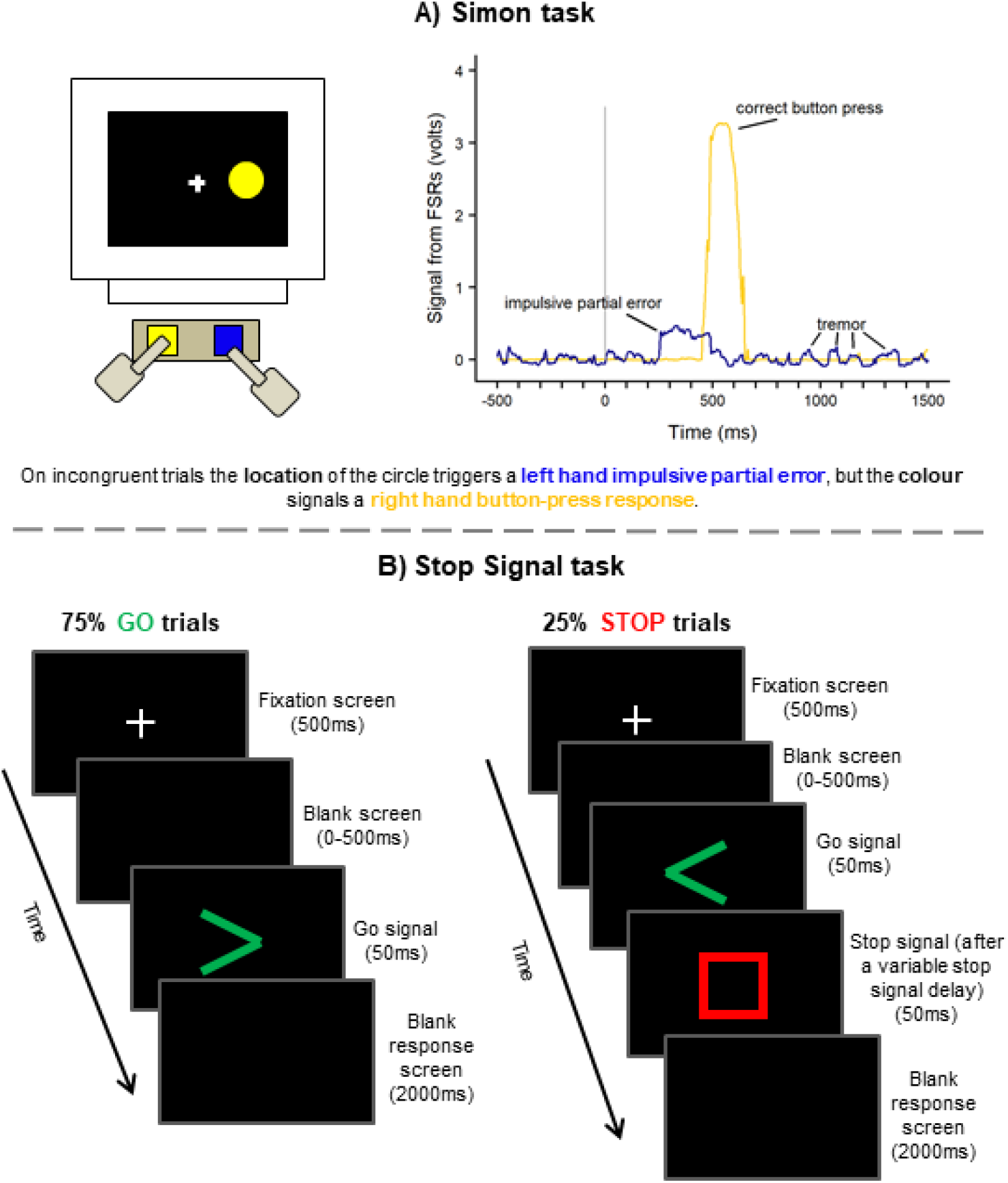
Trial procedure for the Simon task and Stop Signal task. A)In the Simon task, we show an example incongruent trial and the resulting voltage over the course of that trial in a participant with Parkinson’s. The stimulus location (at 0ms) triggered an impulsive right-hand force re-sponse (blue line) that was not detected by the button-press measure. The stimulus colour sig-nalled a left-hand response (yellow) which **was** recorded as a button-press. Data have been smoothed using a 5-point moving average and baseline corrected. In B) Stop Signal task, par-ticipants responded according to the direction of the green arrow, and on 25% of trials attempt to withhold that response upon seeing a red Stop signal after a variable stop signal delay. This delay increases or decreases by 40ms in two 1-up-1-down staircase tracking procedures (independently for each hand) following a successful or unsuccessful Stop trial respectively.

Half of the participants in each group were instructed to press the left key for a blue circle and the right key for a yellow circle, whereas the other half of participants were given the opposite instructions in a counterbalanced design. The stimulus remained on the screen until the participant had made a response, and the next trial began after a 500ms blank inter-trial interval. The experiment consisted of 6 conditions: congruent blue, congruent yellow, incongruent blue, incongruent yellow, neutral blue, and neutral yellow. A trial was said to be “congruent” if the stimulus appeared on the same side of the screen as the side of the response, and “incongruent” if it appeared on the opposite side. In neutral trials the circle was presented centrally. Participants began with a short practice block containing 12 trials (2 trials x 6 conditions). During the practice block, participants were provided with on-screen feedback after each trial (“Correct!” or “Incorrect. Remember, blue = left, and yellow = right” according to counterbalancing) which was not present during the main experiment. The experiment itself consisted of two sessions, approximately an hour apart, each containing four blocks. The first block in each session contained 30 neutral trials (15 of each colour), and the remaining three blocks each contained 80 trials equally split amongst the remaining four conditions. The second session was identical to the first, which resulted in a total of 480 congruent and incongruent trials and 60 neutral trials. Trial order was shuffled randomly and independently for each block and participants were encouraged to rest between blocks

#### Stop Signal task

The Stop Signal task was programmed in Presentation (version 16, http://www.neurobs.com) on the same computer as the Simon task, and using the same method of responding (the left and right keys covered by the FSRs). A white fixation cross (48px) was presented in the centre of a black screen for 500ms, followed by a blank screen for a random duration of 1-500ms to reduce anticipatory responses. The Go signal, a green arrowhead (200 x 200px), was presented in the centre of the screen for 50ms, and participants were instructed to respond with their left or right hand according to the direction of the arrow. On 25% of trials the Stop signal, a hollow red square (250 x 250px), appeared for 50ms after a variable stop signal delay (SSD) which indicated that participants must withhold their pre-potent response to the Go signal (Fig. 1B). The SSD began at 200ms for all participants and was adjusted according to a 1-up-1-down staircase (separately for left and right hands) with a fixed-step of 40ms. Therefore, following a successful Stop (where no button press was recorded) the SSD increased by 40ms on the next stop trial for that hand, and for an incorrect Stop the SSD decreased by 40ms. This procedure helped to ensure that participants were successfully inhibiting their responses on approximately 50% of left and 50% of right-hand Stop trials. Participants were instructed to respond as quickly and accurately as possible and were encouraged not to “wait” to see if a Stop signal would appear (as recommended by Logan, 1994). In both Go and Stop trials, a blank screen was presented after the stimuli for either 2000ms or until a response was recorded.

Participants first completed a practice block consisting of 12 trials during which on-screen feedback was supplied according to the participant’s response (“Correct go”, “Missed button”, “Correct stop”, “Incorrect stop”); this was not present in the main experiment. Participants could repeat the practice until they were comfortable with the task instructions. There were two sessions, approximately an hour apart, each containing 3 blocks. Each block had a total of 120 trials (45 right Go, 45 left Go, 15 right Stop, and 15 left Stop) shuffled randomly and independently for each block. Therefore, there were a total of 720 trials of which 180 were Stop trials.

### Data analysis

Group data were subject to Tukey’s (1977) box plot outlier removal procedure. This removes participants who produced a data point beyond the upper or lower boundaries (3 times the difference between the 25^th^ and 75^th^ percentiles) on any variable within each statistical test.

All data were tested for normality using the Shapiro-Wilk test and then arcsine or log10 transformed (for accuracy and RT data, respectively) if they violated the assumptions of normality. If transformed data still violated the assumptions of normality, then the equivalent non-parametric test was used on the untransformed data. We initially checked to see whether there were differences in performance on both tasks when split by handedness (dominant and non-dominant) but found no significant differences and so collapsed all responses across hands for the remaining analyses.

Alongside null hypothesis significance testing, we calculated Bayes Factors (BF_10_) using default priors in JASP (https://jasp-stats.org/) which demonstrates the likelihood that a particular hypothesis is true given the data. Generally, a BF_10_ below .30 indicates substantial support for the null hypothesis, and a BF_10_ above 3 indicates substantial evidence in favour of the alternative hypothesis (Dienes, 2014; Wagenmakers et al., 2018).

#### Force measurements

Force data were processed using similar methods to those reported in McBride et al. (2012, 2018). In MATLAB R2012a, for each participant and separately for left and right hands, we first smoothed the data using a 5-point moving average; for each data point, an average was taken from that point and the two points either side of it to smooth high frequency noise. The data for each trial were then epoched into 2000ms periods with target onset at 500ms. The first 500ms of the epoch provided baseline activity in the pre-stimulus period which was then used to baseline-correct the following 1500ms on a trial-by-trial basis.

A response was said to have occurred at the first time point in the epoch where the following criteria were satisfied: a recorded amplitude greater than .2 volts^6^ plus 3 standard deviations above the baseline activity, where 17 out of 20 of the following data points also satisfy this criterion, and where another measurement within 70-130% of its amplitude was not detected in the surrounding 250ms. These criteria were chosen^7^ to remain sensitive enough to identify sub-threshold responses that were not forceful enough to produce a button press, whilst remaining conservative enough so as not to erroneously identify instances of tremor from PwP which usually occurs at a frequency of 4-6Hz (Lees, Hardy, & Revesz, 2009). Fig. 1A illustrates a partial, sub-threshold, response in the Simon task from a participant with Parkinson’s who had visible tremor, but where a button-press was recorded in the opposite hand only.

We checked that the force measurement was recording actual button-press responses as expected; full details of this can be found in the supplementary materials.

## Results

We found no reliable interactions between our effects of interest and symptom laterality in PwP (see supplementary materials for full analyses) so the effects of symptom laterality are not reported any further.

### Simon task

#### Button-press data

Accuracy on the Simon task was very high for both groups and in both trial types (accuracy over 96%), so accuracy analyses will not be reported further. Anticipatory RT errors that were likely to have been initiated before stimulus onset (< 150ms) and slow RTs (> 1500ms) were removed first and any remaining outliers removed using Van Selst and Jolicoeur’s (1994) method^8^. One person with Parkinson’s was identified as having very slow overall RTs using Tukey’s (1977) box-plot outlier procedure and was excluded from analysis of such RTs. Summary data and results can be found in Table 2.

**Table 2.** Mean (SD) and statistical tests for the main button-press and response force variables associated with the Simon and Stop Signal tasks in both participant groups.

|  | PwP | HCS | Statistical test |
| --- | --- | --- | --- |
| Congruent RT (ms) | 547 (65) | 543 (58) | $t(45) = .26, p = .80, BF_{10} = .30$ |
| Incongruent RT (ms) | 586 (68) | 583 (63) | $t(45) = .17, p = .86, BF_{10} = .29$ |
| Simon effect for RT (ms) | 39 (23) | 40 (21) | $t(45) = .21, p = .84, BF_{10} = .30$ |
| Congruent partial errors (%) | 9 (5) | 9 (5) | $t(43) = .38, p = .71, BF_{10} = .31$ |
| Incongruent partial errors (%) | 12 (5) | 10 (6) | $t(43) = .84, p = .41, BF_{10} = .39$ |
| Stop accuracy (%) | 55 (4) | 55 (4) | $t(41) = .22, p = .83, BF_{10} = .31$ |
| Go-RT (ms) | 716 (150) | 699 (150) | $t(41) = .36, p = .72, BF_{10} = .32$ |
| SSRT (ms) | 290 (59) | 272 (41) | $t(39.32) = 1.14, p = .26, BF_{10} = .49$ |
| Go partial errors (%) | 10 (5) | 11 (7) | $U = 207, p = .43, BF_{10} = .34$ |
| Stop partial errors (%) | 28 (12) | 27 (14) | $U = 211, p = .38, BF_{10} = .32$ |

A two-way mixed ANOVA showed that for RTs there was a significant main effect of congruency (*F*(1,45) = 153.76, *p* < .001, BF_10_ = 1.084*10^13^) but no significant main effect of group (*F*(1,45) = .05, *p* = .83, BF_10_ = .49) nor an interaction between the effects of congruency and group (*F*(1,45) = .04, *p* = .84, BF_10_ = .28). A raincloud plot of the raw data, median, and interquartile range for RTs can be seen in Fig 2A.

**Fig 2.**
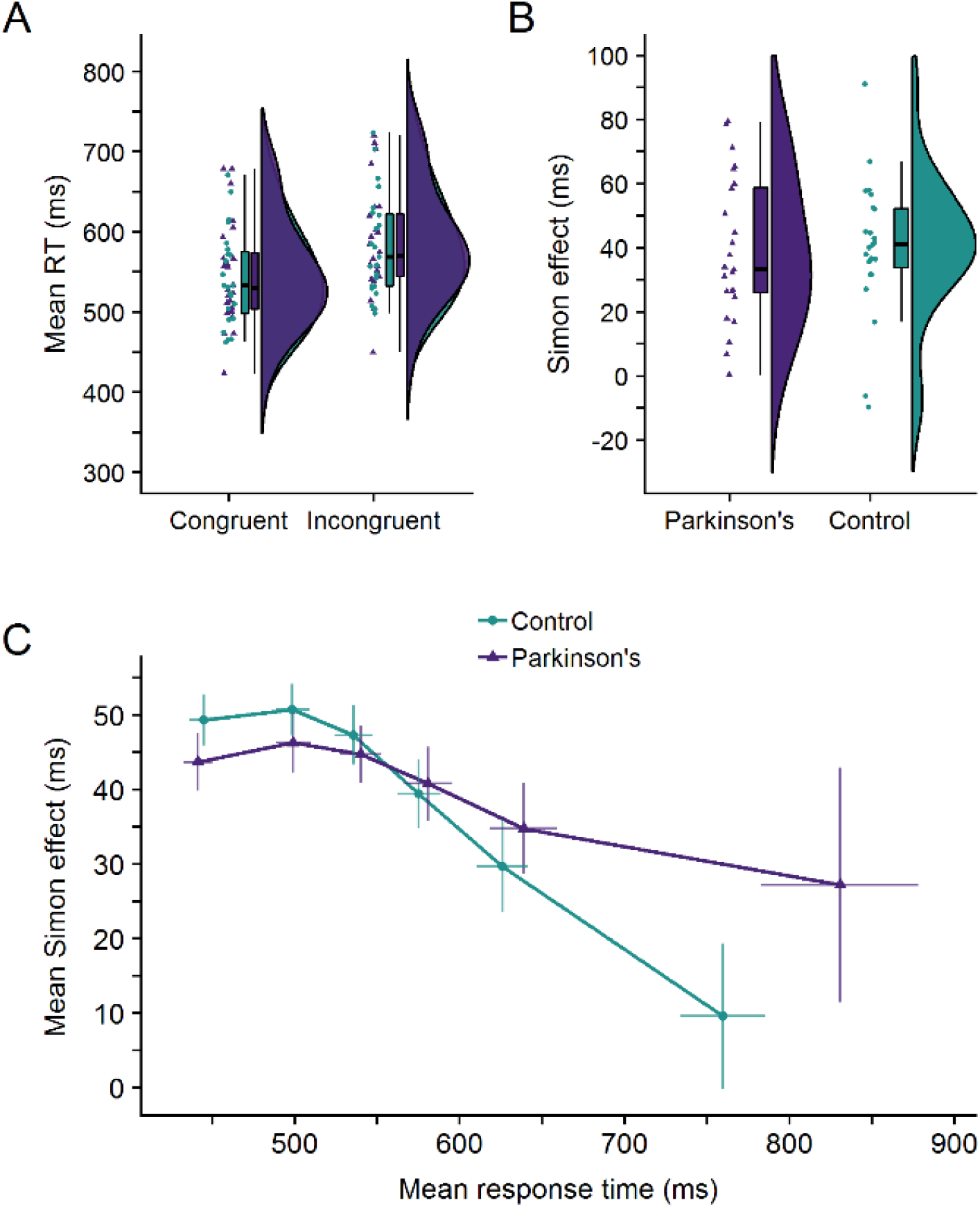
A) A raincloud plot for the response times (RT) in the Simon task on congruent and incongruent trials for both participant groups. The plot displays each participant’s mean correct RT (horizontally jittered), a boxplot, and a split half violin plot of the density (Allen, Poggiali, Whitaker, Marshall, & Kievit, 2018). B) A raincloud plot for the Simon effect (incongruent RT minus congruent RT) for both participant groups. C) Delta plot for the Parkinson’s and healthy control groups. The Simon effect is plotted as a function of RT. Error bars show the standard error of the mean.

#### Distributional analyses

To investigate how the Simon effect changed across the RT distribution, we plotted the Simon effect as a function of the overall correct RT in a delta plot (see e.g, Ridderinkhof, 2002a). Outliers (defined as responses faster than 150ms and slower than 1500ms) and incorrect responses were replaced with the median correct RT for that hand, for that participant, for that condition, with-in that block (to maintain equal bin-sizes). For each participant, RTs were then rank ordered separately for congruent and incongruent trials and divided into 6 equal sized bins (40 trials per bin per condition). The mean RT for each bin in each condition was calculated and then used to calculate the Simon effect (Incongruent RT minus Congruent RT on all correct trials) per bin. The mean Simon effect for each bin was ploted against the mean RT for that bin. The slope between the two bins in the slowest portion of the delta plot is considered the most sensitive measure of response inhibition where a steeper and more negative slope is indicative of greater inhibitory control (Ridderinkhof, 2002a; van den Wildenberg et al., 2010).

Fig. 2C shows the RT distribution for PwP and HCs. A two-way mixed ANOVA showed a significant main effect of slope (*F*(2.78,125.24) = 7.24, *p* < .001, BF_10_ = 1396) but no significant main effect of group (*F*(1,45) = 1.12, *p* = .30, BF_10_ = .30) nor an interaction effect between slope and group (*F*(2.78,125.24) = .33, *p* = .79, BF_10_ = .05). This suggests that whilst susceptibility to the Simon effect does change as a function of RT, as evidenced by a main effect of the gradient of the slopes, this does not differ between PwP and HCs. A planned independent t-test on the gradient of the slope between the slowest two bins additionally revealed that HCs did not have a significantly more negative going final slope compared to PwP (*t*(45) = .65, *p* = .26, BF_10_ = .50, one-tailed).

#### Partial errors in response force

One participant with Parkinson’s and one HC participant were not included in the force analysis for both tasks due to equipment failure on the day of their visit. One further person with Parkinson’s was excluded as an outlier. The data from the FSRs were used to calculate partial errors in response force, that is the percentage of trials containing above-threshold force responses on the incorrect hand where no incorrect button-press was detected. A two-way mixed ANOVA on the percentage of partial errors in response force showed a significant main effect of congruency (F(1,43) = 8.07, p = .007, BF_10_ = 6.17) where more partial errors were detected on incongruent trials compared to congruent trials, but no significant main effect of group (F(1,43) = .46, p = .50, BF_10_ = .45) nor an interaction between the effects of congruency and group (F(1,43) = .37, p = .55, BF_10_ = .33). There were significantly more partial errors on incongruent trials compared to congruent trials for PwP (t(22) = 2.18, p = .02, BF_10_ = 6.78, one-tailed) and HCs (t(21) = 1.86, p = .04, BF_10_ = 1.82, one-tailed), but the Bayes factors suggest the alternative hypothesis is more likely than the null in PwP only. Fig *4A*. shows the raw data, median, and interquartile range for partial errors in response force.

#### Stop Signal task

Five participants were excluded from analysis (2 PwP, 3 HCs) for using a waiting strategy against task instructions. This caused a plateau in the stop signal delays at the maximum available value instead of continually adjusting throughout the task; this left a total of 23 PwP and 20 HCs^9^.

#### Button-press data

Accuracy for Stop trials was expected to be approximately 50% due to the staircase tracking procedure. Go accuracy was very high for both groups (> 97%) so was not analysed further. Anticipatory errors (<150ms) and slow RTs (>1500ms) were removed as outliers, and then any remaining values that were more than 2.5SD away from the mean for each block were also removed. Go-RT was defined as the RT on correct Go trials. There were no significant group differences for any of the above measures (see Table 2).

The SSRT was calculated separately for each hand following the procedure outlined by Verbruggen and Logan (2009): we subtracted the mean SSD from the *N*th percentile of the Go-RT distribution, where *N* is the percentage of failed stops. Although SSRTs were generally longer in PwP (mean = 290ms, SD = 59ms) relative to HCs (mean = 272ms, SD = 41ms), this difference was not significant: *t*(39.32) = 1.14, *p* = .26, BF_10_ = .49. Fig 3. shows the raw data, median, and interquartile range for SSRT.

**Fig 3.**
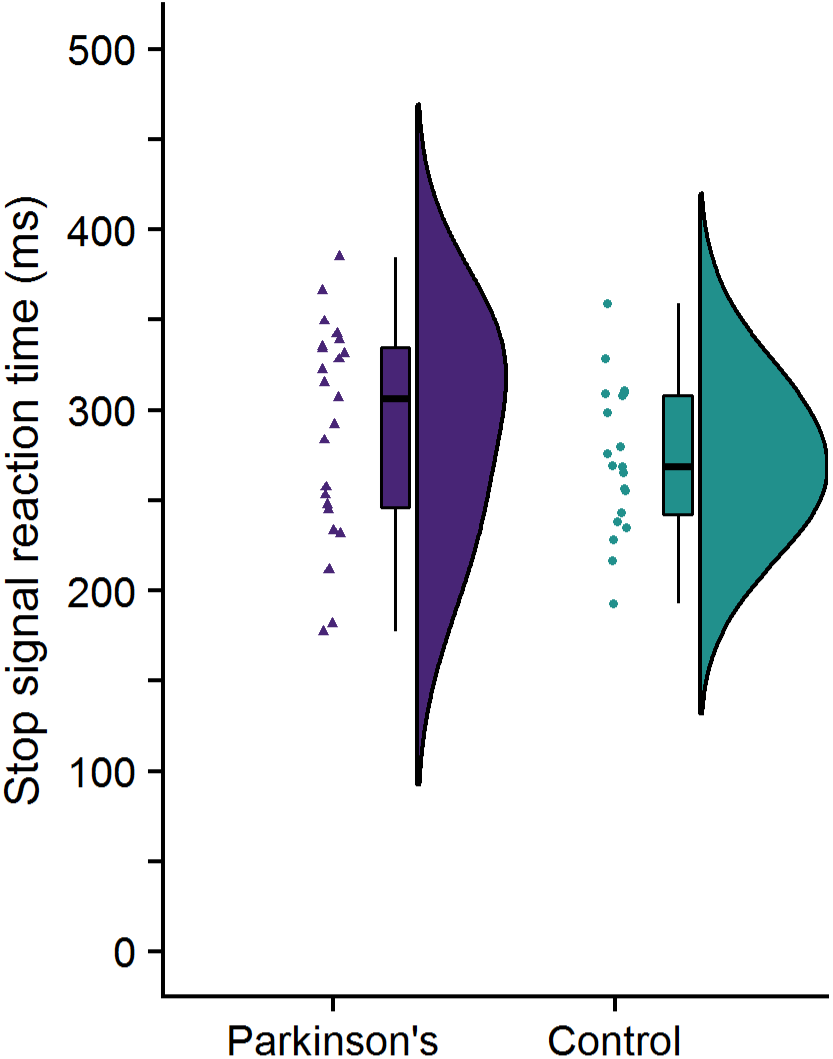
Raincloud plot for the stop signal reaction time (SSRT) for both the Parkinson’s and healthy control groups. The SSRT is an estimation for how long it takes the “Stop” process to overtake the “Go” process for an individual participant.

#### Partial errors in response force

One participant with Parkinson’s was excluded as an outlier. The data from the FSRs were used to calculate partial errors in response force. For Go trials, that is the percentage of trials containing an above-threshold force response on the *incorrect* hand, where a correct button press response had been recorded in the *correct* hand. For Stop trials, this is the percentage of trials that were successfully inhibited according to the button-press data (i.e. no button-press detected), but where an above-threshold force response was detected in the hand *primed to respond* by the direction of the Go signal. Two Mann-Whitney U tests showed that PwP did not produce a significantly higher proportion of partial errors on Go trials compared to HCs (*U* = 207, *p* = .43, BF_10_ = .34, one-tailed) nor on Stop trials (*U* = 211, *p* = .38, BF_10_ = .32, one-tailed). There were significantly more partial errors on Stop trials compared to Go trials for PwP (*t*(20) = 6.15, *p* < .001, BF_10_ = 7932, one-tailed) and HCs (*t*(18) = 5.72, *p* < .001, BF_10_ = 2344, one-tailed). Fig 4B shows the raw data, median, and interquartile range for partial errors in response force.

**Fig 4.**
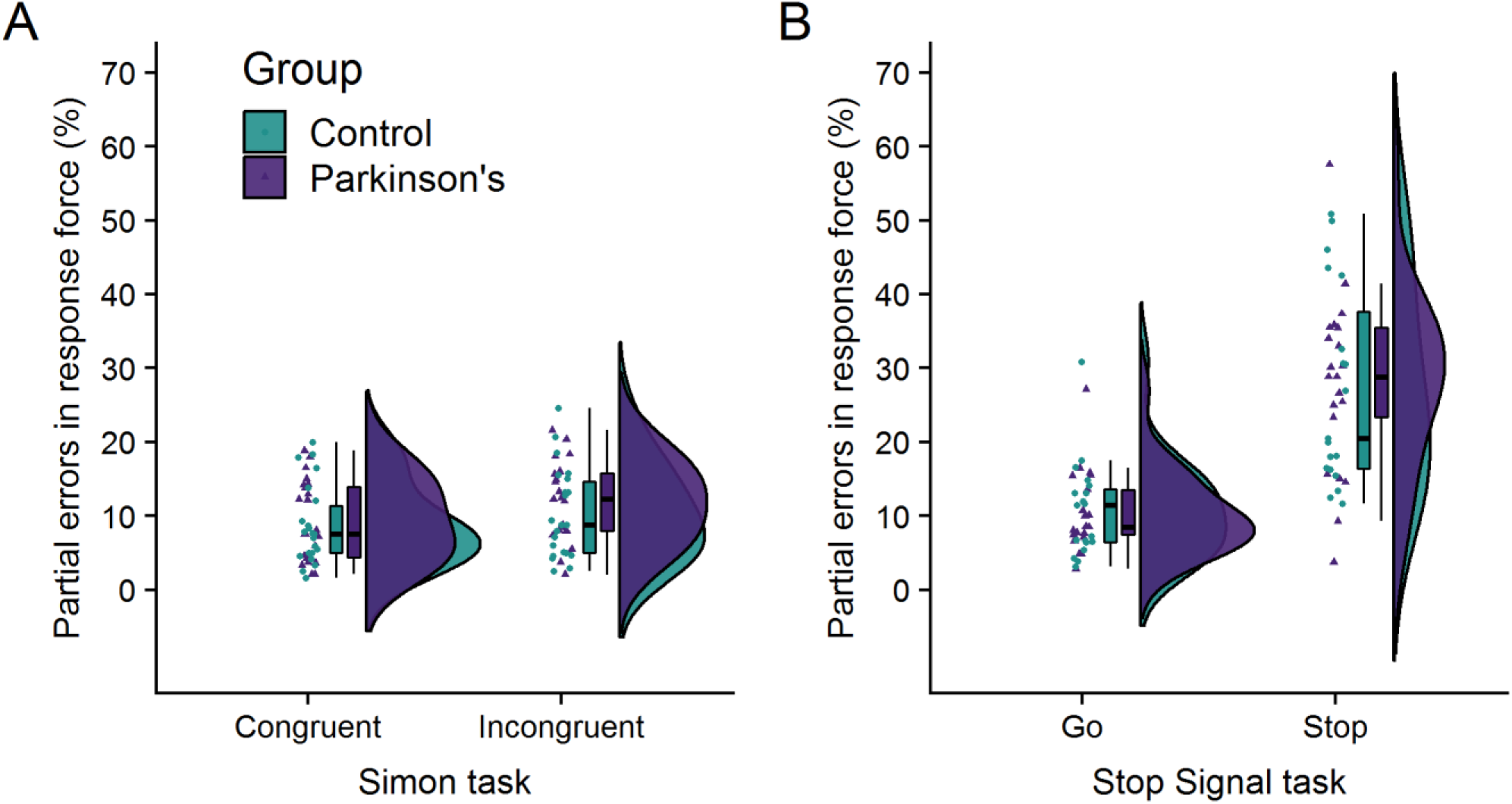
Raincloud plots for partial errors in response force for each group on A) congruent and incongruent trials in the Simon task, and B) Go and Stop trials in the Stop Signal task.

## Discussion

The present study used a continuous measure of response force alongside traditional button-press responses to provide a sensitive behavioural measure of cognitive control in people with Parkinson’s compared to healthy adults in the Simon task and the Stop Signal task. Our button-press data show no significant differences between PwP (at least, with mild-to-moderate symptoms) and HCs with regards to the Simon effect or SSRT, although previous work in this area reports mixed findings. Moreover, and contrary to previous findings reported elsewhere (e.g., van Wouwe et al., 2016; Wylie et al., 2009a, 2009b; Wylie, Ridderinkhof, Bashore et al., 2010; Wylie, Ridderinkhof, Elias et al., 2010), distributional analyses of the time course of our Simon effect showed no significant differences in how well PwP and HCs were able to successfully inhibit responses at the slower end of the RT distribution. As shown in Fig. 2C, the RTs at the slow end of the RT distribution are very variable, particularly for PwP, which may account for the variable findings reported in this field previously.

Such variance may be a feature of any sample of PwP which could suggest that other individual and variable factors of Parkinson’s itself may differentially influence response inhibition.

### Inconclusive group differences in SSRT

The SSRT is an estimation of the time it takes the Stop process to overtake the Go process for each participant. Again, previous research has produced mixed findings. Whilst we found no significant group differences for SSRT, and although our study used a similar number of participants to studies reported elsewhere, our Bayes factors show that we do not have enough evidence to convincingly accept or reject the null hypothesis). Potentially, this explains the mixed findings in the literature thus far; many studies are underpowered (Dumas-Mallet, Button, Boraud, Gonon, & Munafò, 2017) and there may not yet be enough evidence in the literature to conclude whether PwP have difficulties with response inhibition.

### Both groups show more partial errors in response force for trials requiring response inhibition

Partial errors in response force on incongruent trials may reflect the cognitive process of suppressing an automatically activated response in favour of the goal directed response (Ridderinkhof, 2002a, 2002b; van den Wildenberg et al., 2010). We sought to use this measure to complement previous research that detected a functional deficit in PwP even where no behavioural deficit was present (Baglio et al, 2011; Vriend et al., 2015). We used these data to detect partial errors in response force; that is, where an increase in response force is detected either in the absence of a button-press (on Stop trials in the Stop Signal task), or where a button-press was detected in the *opposite* hand (in the Simon task, and on Go trials for the Stop Signal task). On the Simon task, both groups made significantly more partial errors in response force for incongruent trials compared to congruent trials. There were no group differences which may suggest that there is no functional deficit present in Parkinson’s if our response force measure is sensitive enough to pick up more subtle differences in response conflict. Interestingly, the Bayes factors suggest that there is more evidence for the conclusion that PwP produce more partial errors on incongruent than congruent trials, but that in HC participants there is insufficient evidence to support the statistically significant difference and to confidently reject the null hypothesis. This could be tentatively interpreted in opposing ways. Firstly, this may reflect *better* response inhibition in PwP as they may be better able to suppress the response before it produces an incorrect button-press, whereas in HCs these partial responses may be more likely to result in an incorrect button-press. Alternatively, it could reflect *worse* response inhibition in PwP. HCs may be able to suppress their responses faster and produce fewer partial errors in response force for this reason, as the suppression successfully occurs earlier in the potential motor movement.

On the Stop Signal task, partial errors (where there was above-threshold force applied to the response button, but this force was not sufficient for a button-press to be detected) were recorded on up to 30% of Stop trials which demonstrates that our measure provides a sensitive means of detecting sub-threshold erroneous response activity in the effectors that would otherwise be missed by conventional button-press measures alone. Moreover, there was no significant difference in the number of partial errors recorded for PwP compared to healthy controls, and indeed our Bayes factors indicate that partial error rate was perhaps equivalent for the two groups (BF_10_ = .32).

### Does performance on different tasks correlate?

The Simon effect was significantly and positively correlated with the total Barratt Impulsiveness Score, but not the motor score (see supplementary materials). Therefore, a higher score of trait impulsivity is correlated with a larger Simon effect. This finding is consistent with previous research from Duprez et al. (2017); they found significant correlations between total impulsivity score and increased impulsive errors. However, they also found that total impulsivity is also correlated with *better* inhibitory control at the slow end of the RT distribution; they suggest the subthalamic nucleus, part of the basal ganglia circuitry affected in Parkinson’s, has a temporally dissociated role in both poor conflict resolution and successful response suppression, as well as involvement in trait impulsivity (Duprez et al., 2017) which may help explain our correlation here.

The SSRT did not correlate significantly with the Barratt Impulsiveness Scale total or motor scores which suggests that trait impulsivity, especially when related to motor impulsivity (BF_10_ = .45), is unrelated to the ability to withhold a response, contrary to previous findings (Caswell, Bond, Duka, & Morgan, 2015; Gorlyn, Keilp, Tryon, & Mann, 2005; Nolan, D’Angelo, & Hoptman, 2011). Previous work has also suggested that the factor structure of the Barratt Impulsiveness Scale might be different in PwP compared to the general population, as there is low internal consistency (Smulders, Esselink, Cools, & Bloem, 2014), and indeed a different factor structure does appear to exist in PwP (Ahearn, McDonald, Barraclough, & Leroi, 2012).

We also found no significant correlation between the Simon effect and SSRT for PwP. Although previous research has suggested that there is an overlap in the brain networks required to perform successfully in both tasks (Jahfari et al., 2011; Sebastian et al., 2013), our data may suggest that the tasks load different mechanisms of inhibition and control.

### Limitations of the current study

Parkinson’s is a heterogeneous disease and, as such, it is difficult to compare samples across studies. Generally, participants with Parkinson’s tend to have more mild symptoms, owing to the practicalities of needing to be able to perform the task(s) (e.g. make a response using a button-box) which limits the generalisability of any findings to more advanced Parkinson’s cases. PwP across studies often exhibit a mix of confounding characteristics, some of which have been shown to affect response inhibition and response conflict in other studies, such as subthalamic nucleus deep brain stimulation and the presence of additional ICBs (Mirabella et al., 2012; Ray et al., 2009; Swann et al., 2011; van Wouwe et al., 2016; van den Wildenberg et al., 2006; Wylie, Ridderinkhof, Bashore et al., 2010; Wylie, Ridderinkhof, Elias et al., 2010; Wylie et al., 2012). Whilst there were no participants with deep brain stimulation in our present sample, much of the literature - including this study where we did not collect such information - do not specifically exclude or account for PwP who have additional ICBs, which may well be up to 50% of any sample (Corvol et al., 2018). It is therefore likely that an unknown proportion of any sample of participants with Parkinson’s also have ICBs, which will affect any conclusions made about the effects of Parkinson’s (relative to dopaminergic medication) on response inhibition.

It is also possible that the published literature may overestimate group differences. As noted above, we have used similar tasks with a similar number of participants to many of the studies reported in the literature, and yet our Bayes factors indicate that we do not always have enough evidence to accept or reject the null hypothesis. In those cases where we did have enough evidence, it was largely in favour of the null hypothesis that there are no significant differences in response inhibition between PwP and HCs.

The force response analysis used here was built upon previous work by McBride et al. (2012, 2013, 2018), and specifically adapted to be suitable for PwP. The data from the PwP had a lower signal-to-noise ratio than data from the HCs due to many PwP exhibiting the tremor that is often associated with their disease. We attempted to account for this during data analysis by filtering out above-threshold responses that occurred at a frequency of a typical Parkinsonian tremor (4-6Hz, Lees et al., 2009). It is therefore possible we are missing some genuine responses or mistakenly categorising tremor or random noise as a genuine response. Despite these possible imperfections, this measure still provides a more sensitive measure than button-presses, as shown by our ability to capture partial errors in response force that were not detected in the button-presses.

It is additionally possible that we may be mistakenly categorising mirror movements as partial errors in response force. Mirror movements are simultaneous movements of a lesser amplitude that can occur in the opposite hand to the one performing an action and were observed here visually early in the analysis process. They tend to be pathological in nature after childhood and are particularly prominent in the earlier stages of Parkinson’s (Beaulé et al., 2012; Espay, Li, Johnston, Chen, & Lang, 2005). We cannot assume that mirror movements occur independently of response inhibition and may therefore be unequally distributed across trials requiring, or not requiring, an inhibitory process. With our current method it is difficult to define and distinguish mirror movements from partial errors of inhibitory control.

Our algorithm was designed to remove the effects of Parkinson’s tremor from the force data (so tremors were not mistaken for partial responses), and this necessarily means that we have not analysed any effects of tremor phase on participants’ performance on the tasks presented here. It is possible that errors may have been elicited more commonly during different phases of the participants’ tremor – such as when the stimulus onset as the tremor was in the same direction as the response. We would expect any such effect of stimulus presentation coinciding with tremor phase to be equally distributed across conditions and so is unlikely to account for any effects reported here, but this might be a fruitful avenue for further investigation.

## Conclusions

Overall, we provide evidence that PwP and HCs do not significantly differ on their susceptibility to the Simon effect using button-press measures, but insufficient evidence regarding group differences for the percentage of partial errors on incongruent trials in this task. Conversely, we found insufficient evidence to support the null hypothesis that SSRTs do not differ between groups, but evidence in favour of the null hypothesis that the groups produce a similar percentage of partial errors in response force on Stop trials. In summary, we show that it is more likely that people with mild-to-moderate Parkinson’s do not show an impairment in response inhibition or response conflict relative to healthy controls, but that more evidence is needed to make even stronger conclusions in favour of the null.

Additionally, we demonstrated the utility of a more sensitive method of measuring the cognitive process of response inhibition and response conflict using force sensing resistors; this allowed us to identify partial responses that would have gone undetected by conventional button-press measures (including up to 30% of trials in the Stop Signal task). This may be a useful tool to detect more subtle group differences in tasks of ongoing cognitive control that are usually measured with button-press responses.

## Footnotes

1 Data were collected from one additional participant with Parkinson’s, but the severity of their tremor meant they were not able to satisfactorily complete the tasks. Their data were not analysed.

2 An anonymous reviewer suggested we check there were no differences in impulsivity on all main variables in the experimental analyses for patients on dopamine agonists medication vs those without. For all analyses (including BIS scores), there were no differences between these patient groups.

3 During data collection, we discovered that two items from the Barratt Impulsiveness Scale (“I plan for job security”, nonplanning impulsivity; “I change jobs”, motor impulsivity) were often irrelevant for this largely retired demographic, and after data collection we confirmed that this comprised the majority of the missing data. These items were therefore removed from analysis for all participants and any remaining missing data were replaced with the mean score for that sub-scale for that participant.

4 As suggested by an anonymous reviewer, we repeated the main analyses in an exploratory manner with age and GDS scores as covari-ates to check that the significant group differences on these measures did not affect the pattern of results. We found that our conclusions remained the same.

5 The MDS-UPDRS was used to identify tremor dominant and postural instability and gait dominant patients using the same method re-ported by Stebbins et al. (2013).

6 As in McBride et al. (2018) we used a constant in addition to a standard deviation threshold in order to reject noise and more reliably detect responses.

7 The researchers were blind to the condition and group when making decisions as to how to process the data.

8 This method trims outliers with a per condition and per participant moving standard deviation, where the standard deviation is adapted depending on the number of trials

9 After removal of these participants’ data there were no other meaningful changes to group differences on demographic or neuropsychological measures.

