## Supplementary materials for "Continuous force measurements reveal no inhibitory control deficits in Parkinson’s disease"

### 1. Correspondence between PwP and HC’s button-presses and response force measures

As with any threshold, it is possible that we are missing some genuine responses in the force measurement and/or erroneously categorising some random noise as genuine response – the same would be true of any measure. To determine whether there are any systematic effects of the threshold we have chosen we examined what percentage of button press responses were also detected as being above force threshold, i.e. the “correspondence” between the two measures.

Overall, and reassuringly, correspondence was high for both tasks (see Table S1). This means that in most trials where a binary button-press was detected, there was also an identifiable response detected by the algorithm for the force sensing resistors (FSRs) data. There was a significantly higher correspondence for the HCs relative to the PwP (a mean of 98-99% correspondence compared to a mean of 95-96% correspondence), which most likely reflects the Parkinsonian tremor making it more difficult for our algorithm to discriminate tremor from response with 100% accuracy. Although this is a small difference in real terms, it might mean that our response force measure may miss more responses in PwP than in the HC group. Despite this, response force is still more sensitive than the standard button press measurement.

**Table S1. The correspondence is the percentage of trials with a button-press where a response was also detected by the algorithm on the data from the FSRs**

| | PwP | HCs | Statistical test |
|---|---|---|---|
| Simon - Congruent | 95 (8) | 99 (1) | $U$ = 126, $p$ = .004, $BF_{10}$ = 7.97 |
| Simon - Incongruent | 95 (6) | 98 (1) | $U$ = 158, $p$ = .03, $BF_{10}$ = 1.85 |
| SST – Go | 96 (4) | 98 (2) | $U$ = 120, $p$ = .03, $BF_{10}$ = 1.38 |

### 2. Symptom laterality

#### 2.1 Simon task

A two-way within-subjects ANOVA with a factor of congruency (congruent, incongruent) and laterality (more-affected hand, less-affected hand) on RT showed a main effect of congruency ($F(1,23) = 72$, $p < .001$, $BF_{10} = .16$), but no main effect of laterality ($F(1,23) = 4.18$, $p = 0.05$, $BF_{10} = 0$) nor an interaction effect between laterality and congruency ($F(1,23) = 0$, $p > 0.99$, $BF_{10} = 0$).

A two-way within-subjects ANOVA on the percentage of partial errors in response force in the Parkinson’s participants showed a significant main effect of congruency ($F(1,22) = 4.75$, $p = .04$, $BF_{10} = 2.71$) but no main effect of symptom laterality ($F(1,22) = .03$, $p = .88$, $BF_{10} =0.22$), and no interaction effect between congruency and symptom laterality ($F(1,22) = 1.44$, $p = .24$, $BF_{10} = .32$).

#### 2.2 Stop Signal Task

**Table S2. Data for the Stop Signal task by more-affected and less-affected hand in people with Parkinson's**

| | More-affected hand | Less-affected hand | Statistical test |
|---|---|---|---|
| Go RT | 727ms ± 155ms | 705ms ± 147ms | $t(22) = 3.08$, $p = .006$, $BF_{10} = 8.13$ |
| SSRT | 292ms ± 60ms | 287ms ± 64ms | $t(22) = .78$, $p = .45$, $BF_{10} = .29$ |
| Partial errors - Go | 12% ± 6% | 9% ± 6% | $t(20) = 1.89$, $p = .07$, $BF_{10} = 1.01$ |
| Partial errors - Stop | 29% ± 15% | 27% ± 13% | $t(20) = .54$, $p = .60$, $BF_{10} = .26$ |

### 3. Correlations between response inhibition, response conflict, and trait impulsivity

As an exploratory addition, we also examined trait impulsivity with the Barratt Impulsiveness Scale (BIS; Barratt, 1959; Patton, Stanford, & Barratt, 1995), which has been shown to be generally higher in PwP compared to HCs (Isaias et al., 2008; Nombela et al., 2014).

We performed exploratory correlations to examine the relationship between SSRT (response inhibition), Simon effect (response conflict), and the Barratt Impulsiveness Scale's total and motor impulsivity scores (Patton et al., 1995). Exploratory Pearson's *r* correlations (Bonferroni corrected, $a = .01$) were performed in each group.

**Table S3. Pearson's correlations for people with Parkinson's and the healthy control group**

| | | Parkinson's | | | Healthy controls | | |
|---|---|---|---|---|---|---|---|
| | | Pearson's *r* | *p* | $BF_{10}$ | Pearson's *r* | *p* | $BF_{10}$ |
| BIS (total) | SSRT | .26 | .11 | .90 | -.24 | .85 | .15 |
| BIS (total) | Simon effect | .50 | .006* | 9.86 | .17 | .22 | .53 |
| BIS (motor) | SSRT | .14 | .27 | .45 | -.22 | .83 | .15 |
| BIS (motor) | Simon effect | .27 | .10 | .98 | .04 | .42 | .30 |
| SSRT | Simon effect | .41 | .03 | 2.80 | .29 | .11 | .96 |

As shown in Table. S3, there was a significant positive correlation between the size of the Simon effect and the total impulsiveness score for people with Parkinson's at the adjusted *a* level ($r = 0.50$, $p = .006$, $BF_{10} = 9.86$), but not the healthy controls. This suggests some problems with impulsivity for PwP which may affect response conflict and trait impulsivity, but not response inhibition as measured with SSRT. There are higher impulsiveness scores and greater variability in scores for the Parkinson's group and not the HC group which may contribute to some findings; it could be that this applies to a subset of the patients such as those with undiagnosed/unreported impulse control disorders.

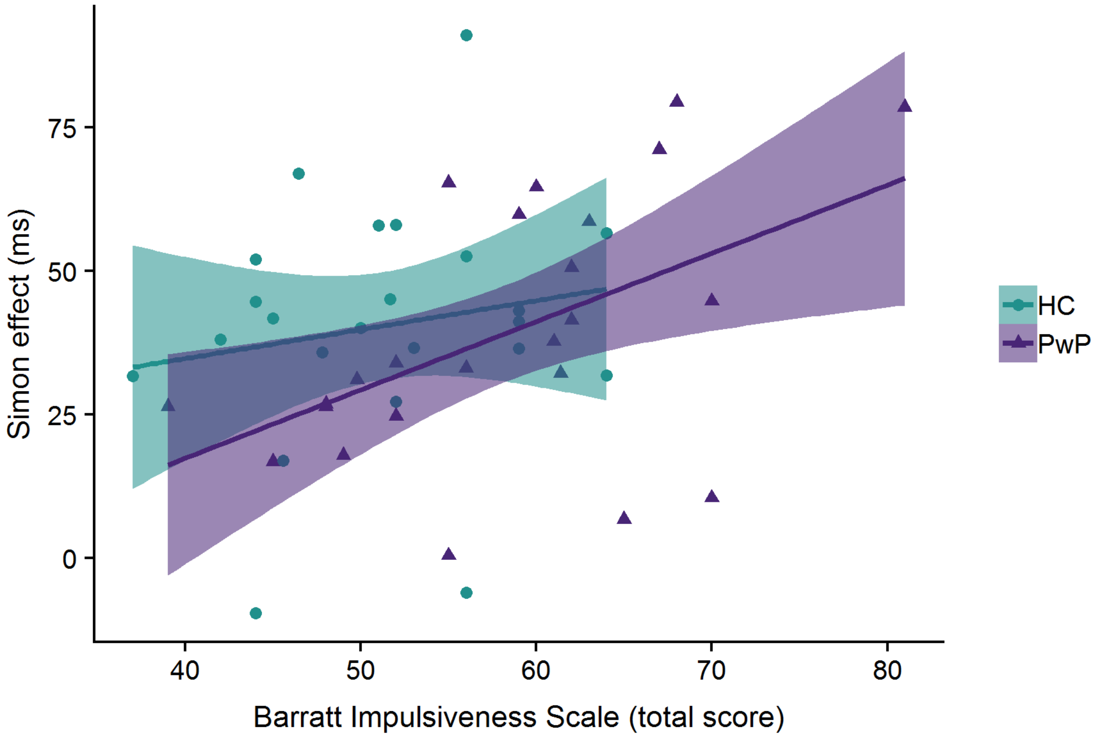


**Fig S1. Scatterplot of the correlation between the Simon effect and total score on the Barratt Impulsiveness Scale for PwP and HCs. The data is shown with a linear regression line of best fit and shaded confidence intervals.**

There were no other significant correlations between any of the measures taken for either participant group.
